# Thalamocortical axons regulate neurogenesis and laminar fates in early sensory cortex

**DOI:** 10.1101/2021.06.16.448668

**Authors:** Timothy Monko, Jaclyn Rebertus, Jeff Stolley, Stephen R. Salton, Yasushi Nakagawa

## Abstract

Area-specific axonal projections from the mammalian thalamus shape unique cellular organization in target areas in the adult neocortex. How these axons control neurogenesis and early neuronal fate specification is poorly understood. By using mutant mice lacking the majority of thalamocortical axons, we show that these axons increase the number of layer 4 neurons in primary sensory areas by enhancing neurogenesis and shifting the fate of superficial layer neurons to that of layer 4 by the neonatal stage. Part of these area-specific roles is played by the thalamus-derived molecule, VGF. Our work reveals that extrinsic cues from sensory thalamic projections have an early role in the formation of cortical cytoarchitecture by enhancing the production and specification of layer 4 neurons.

## Introduction

The adult mammalian nervous system has a remarkable complexity with thousands of neuronal types each of which adopts consistent numbers and locations. This fundamental feature critically depends on early developmental mechanisms that control cell divisions and cell fate specification. A unique feature of the nervous system is that neurons can grow axons over great distances and interact with cells far from their cell body. Although such interactions are known to control survival and maturation of the target neurons, they could also influence both the absolute and relative abundance of each neuronal type in the target region. However, we currently have little knowledge on such roles of afferent axonal projections. The mammalian neocortex is composed of diverse types of neurons that are sequentially generated during embryonic development and are arranged in an orderly manner across layers (McConnell, 1989; Gao et al., 2014; Llorca et al., 2019; Oberst et al., 2019). In adult neocortex, the distribution and density of each neuronal type differ between areas. Primary (first-order) visual, somatosensory and auditory areas have well-developed layer 4 that is directly targeted by axons from corresponding principal sensory nuclei of the thalamus (Brodmann, 1909; Peters and Jones, 1984). Early patterning mechanisms within the cortex intrinsically confer positional identity of neural progenitor cells, which leads to the formation of area maps in postnatal cortex (Rakic, 1988; Bishop et al., 2000; Fukuchi-Shimogori and Grove, 2001, 2003). These mechanisms influence region-specific regulation of neurogenesis and subsequent specification of neuronal types, which will eventually control the formation of area-specific cytoarchitecture and functional specialization of the neocortex. In contrast, topographic and area-specific projections of thalamocortical axons (TCAs) provide a major extrinsic influence on neocortical cytoarchitecture (Rakic, 1988; O’Leary, 1989; López-Bendito, 2018; Cadwell et al., 2019; Nakagawa, 2019). For example, ablation of principal sensory thalamic nuclei or blocking transmitter release from TCA terminals during the postnatal period disrupts anatomical and molecular hallmarks of the primary sensory cortex (Wise and Jones, 1978; Narboux-Nême et al., 2012; Chou et al., 2013; Li et al., 2013; Pouchelon et al., 2014). In addition, disrupting the intensity and temporal pattern of spontaneous activity of embryonic thalamic neurons leads to the lack of normal anatomical and functional organization of the primary somatosensory cortex (Antón-Bolaños et al., 2019). Because TCAs arrive in the cortex during the early period of superficial layer (layers 2-4) neurogenesis (Fig.1A,B), production and fate specification of these neurons could be modified by TCAs during embryogenesis. Although eye removal experiments in primate and carnivore visual systems suggested the roles of the thalamic input in cortical neurogenesis (Dehay et al., 1991; Reillo et al., 2017), how TCAs modulate generation and specification of neocortical neurons in an area-specific manner has remained elusive. In this study, we used a mouse model in which embryonic TCA development is disrupted, and found evidence indicating that TCAs not only enhance the numbers of progenitor cells but also promote the layer 4 fate of newly generated superficial layer neurons at the expense of the layer 2/3 fate. This early influence of TCAs, combined with their later roles in activity-dependent neuronal maturation, contribute to the characteristic cytoarchitecture of primary sensory cortex in the adult brain. We further provide evidence that the TCA-derived molecule VGF confers sensory-specific regulation of progenitor cell division and neonatal cell fate specification.

**Figure 1.**
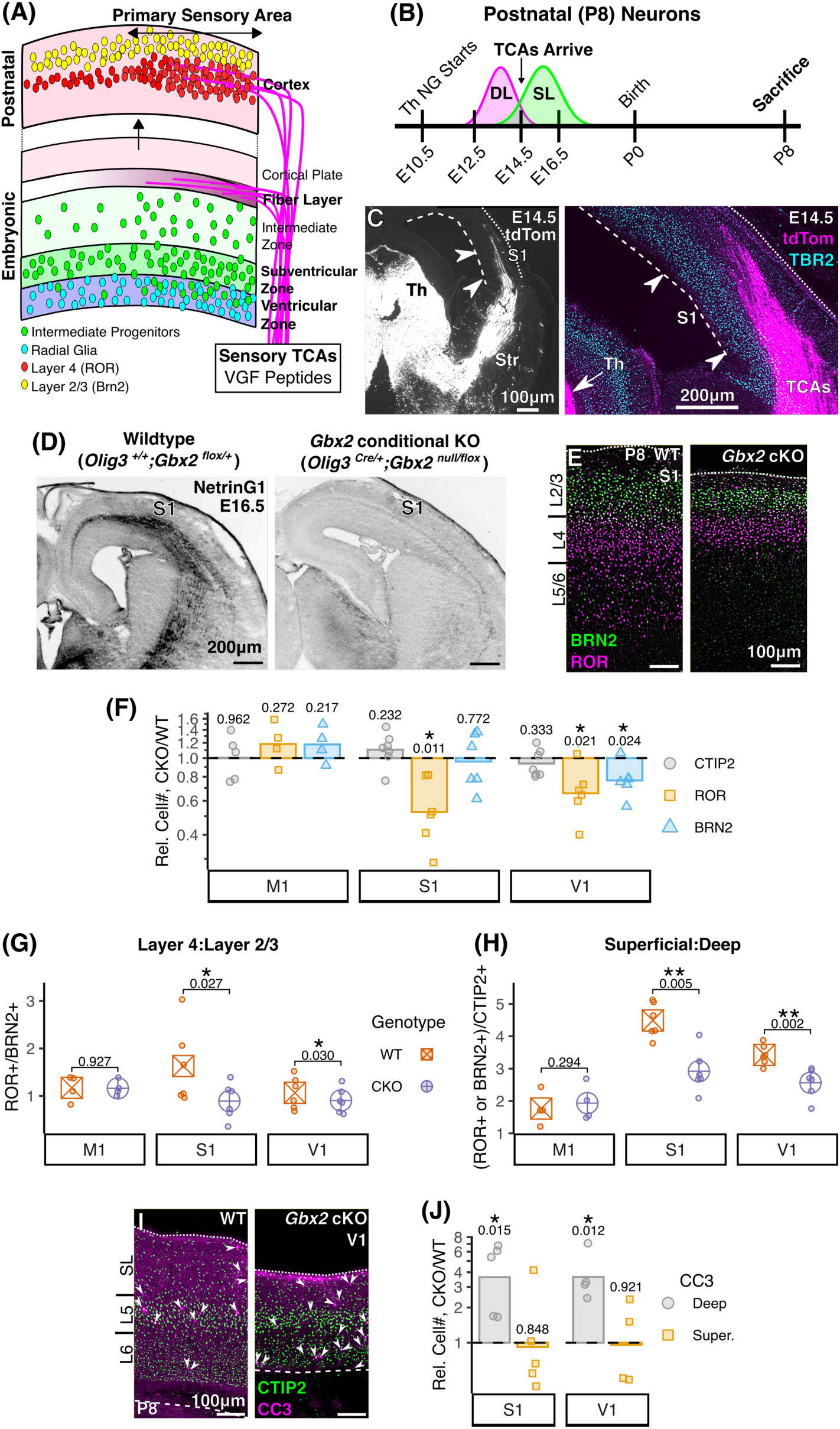
Thalamocortical axons increase postnatal neuron number in primary sensory cortex. **(A)** Schematic summary of our findings; TCAs, including sensory-specific VGF, increase the production of superficial layer neurons and bias their fates towards layer 4. **(B)** Experimental scheme in relation to the timing of neocortical neurogenesis (DL: deep layer; SL: superficial layer) and arrival of TCAs; Th NG: thalamic neurogenesis. **(C)** TCAs (tdTomato driven by *Olig3^Cre^*) project to cortex and reside near TBR2^+^ progenitors in E14.5 S1 cortex. **(D)** Comparison of wildtype (WT) and *Gbx2* conditional knockout (cKO) thalamocortical axons (TCAs) labeled by NetrinG1 (inverted). At E16.5, NetrinG1 is found throughout the cortex in WT mice, but in *Gbx2* cKO mice few reach the cortex. **(E)** Layer 4 (ROR^+^) and layers 2/3 (BRN2^+^) neurons in S1 cortex in wildtype (WT) and *Gbx2* cKO mice. **(F)** SL neurons are decreased in both S1 (ROR) and V1 (ROR & BRN2) in *Gbx2* cKO mice; DL (CTIP2) unchanged. **(G-H)** S1 and V1 cortex in *Gbx2* cKO mice have decreased ratios of ROR^+^-to-BRN2^+^ neurons and SL-to-DL neurons compared with WT controls. **(I)** CTIP2 (DL) and cleaved caspase 3 (CC3: apoptotic cells) in V1 at P8. **(J)** CC3^+^ cells are increased within DLs (bounded by CTIP2^+^ cells) of *Gbx2* cKO mice. Matched ratio t-test (F,J) or paired t-test (G,H) *p*-values shown between WT and *Gbx2* cKO. For (F,J), data shown as relative difference of *Gbx2* cKO to WT littermates (dashed line); points represent individual pairs, bar represents pooled mean.

## Results

### Thalamocortical axons enhance the numbers of superficial layer neurons in primary sensory cortex

By embryonic day 14.5 (E14.5) in mice, thalamocortical axons (TCAs) have already reached the prospective somatosensory area and spatially overlap with the domain occupied by intermediate progenitor cells (IPCs) (Fig.1C, S1A) and radial glia fibers that span from the ventricular lumen to the pial surface. This timing coincides with the onset of superficial layer neurogenesis in this area (Fig.1B)(Klingler et al., 2019). In the prospective visual cortex, TCAs have just reached its lateral portion by this stage (Fig.S1A,B). Newly generated superficial layer neurons in embryonic neocortex lack patterns of gene expression, morphology and connectivity that distinguish precise laminar identity in the postnatal brain (Oishi et al., 2016; De León Reyes et al., 2019; Di Bella et al., 2020). Prospective layer 4 neurons start to show characteristic gene expression by E18.5 (Di Bella et al., 2020), and become spatially segregated from layer 2/3 neurons by postnatal day 1 (P1) (Oishi et al., 2016). Between E14.5 and P1, TCAs are in close proximity to both neural progenitor cells and immature neurons (Fig.S1C-F). Therefore, TCAs could interact with these cells and control the generation and/or specification of superficial layer neurons before direct thalamocortical synapses are formed (Agmon et al., 1996; Molnár et al., 2003). In order to understand the early roles of TCAs, we analyzed mutant mice (thalamus-specific, *Gbx2* conditional knockout mice; *Gbx2* cKO mice, *Olig3^Cre/+^; Gbx2 ^null/flox^*) in which most TCAs fail to project to the cortex (Fig.1D,S1A)(Vue et al., 2013). Postnatal viability of the *Gbx2* cKO mice allowed us to assess the full extent of the TCAs’ roles in neocortical development starting at the mid-embryonic into postnatal stages. At P8, *Gbx2* cKO mice showed a thinning of layer 4 and blurring of the borders of primary sensory areas (Fig.S1G) (Vue et al., 2013). To determine if neuron numbers are altered in the mutant cortex, we performed immunohistochemistry for transcription factors that are essential for functional identify of representative classes of excitatory neocortical neurons, including RORα/β (layer 4; with anti-ROR antibody, Fig.S1H,I), BRN1/2 (POU3F1/2) (layer 2/3; with anti-BRN2 antibody (Yamanaka et al., 2010)), and CTIP2 (BCL11B) (layer 5/6) (Sugitani et al., 2002; Arlotta et al., 2005; Chen et al., 2008; Jabaudon et al., 2011; Dominguez et al., 2013; Clark et al., 2020) (Fig.S1J). Among these, RORβ is both necessary and sufficient for the acquisition of characteristic morphology and gene expression patterns of layer 4 spiny stellate neurons (Jabaudon et al., 2011; Clark et al., 2020). We compared the numbers of positive cells between brains of cKO and wildtype littermates with unbiased threshold-based segmentation, and found that the lack of TCAs in *Gbx2* cKO cortex results in significantly decreased numbers of ROR-positive (ROR^+^) layer 4 neurons compared with wildtype controls in both primary somatosensory (S1) and visual (V1) areas, but not in primary motor area (M1) (Fig.1E,F, Fig.S1K,L). The number of layer 2/3 neurons was also decreased in V1, while layer 5 neurons were not decreased in any of the areas analyzed (Fig.1F). In both S1 and V1 but not in M1, the ratio of layer 4 (ROR^+^) cells to layer 2/3 (BRN2^+^) cells was decreased in the cKO cortex (Fig.1G), as was the ratio of superficial (ROR^+^ and BRN2^+^) to deep (CTIP2^+^) layer neurons (Fig.1H). Thus by P8, TCAs normally enhance the number of superficial layer neurons as well as the ratio of layer 4 neurons to layer 2/3 neurons in primary sensory cortex but not in motor cortex. Because TCAs could provide a survival signal for neurons in their target region, we counted apoptotic cells at P8 and found no increase in cleaved caspase 3 (CC3)-positive cells in superficial layers of *Gbx2 cKO* mice (Fig.1I,J). Instead, *Gbx2 cKO* mice had increased apoptotic cells in deep layers. This may be due to the decreased corticofugal connectivity (Hevner et al., 2002) and the absence of retrograde survival factors provided by downstream target areas, including the thalamus, in the absence of TCA projections.

### Thalamocortical axons increase progenitor cells in embryonic sensory cortex

The above results prompted us to determine if the decrease in superficial layer neurons in early postnatal *Gbx2* cKO mice is due to their decreased production from progenitor cells during embryogenesis. At E16.5, *Gbx2* cKO mice had reduced numbers of progenitors towards the end of superficial layer neurogenesis (Fig.2A); the number of radial glia (RGs; PAX6^+^) was decreased in prospective S1 and intermediate progenitor cells (IPCs; TBR2^+^) were also decreased in both prospective S1 and V1, but no changes were observed in prospective M1 and medial prefrontal cortex (Fig.2B-E, S2A-D). The number of IPCs was already decreased in prospective S1 and V1 at E14.5 (Fig.2G-H, S2E-F) when these cells mainly produce layer 4 neurons (Klingler et al., 2019) (Fig.2A). Mitotic cells labeled by an antibody against phosphorylated histone H3 (PH3) were also decreased at both basal and apical locations at E16.5 in prospective S1 and V1 (Fig.2E-F), as well as at E14.5 in prospective S1 (Fig.2I). Thus, TCAs appear to increase the number of progenitors and their divisions during the production of superficial layer neurons. To test whether progenitor cells show an increased tendency to exit the cell cycle in the absence of TCAs, which would result in loss of progenitors, we pulse-labeled S-phase cells with ethynyldeoxyuridine (EdU) 18 hours prior to analysis at E16.5 (Fig.2J), which allowed us to investigate the short-term fate of daughter cells of the labeled progenitors (Takahashi et al., 1995; Wang et al., 2011). Both the number of Ki67-positive, proliferating cells and EdU-labeled cells were decreased in both prospective S1 and V1 in cKO mice (Fig.2K). The rate of cell cycle re-entry as measured by the Ki67^+^EdU^+^/EdU^+^ ratio was also decreased (Fig.2L-M, S2G-H). These results collectively demonstrate that in primary sensory cortex, TCAs contribute to increased production of superficial layer neurons by enhancing the number of progenitor cells and prolonging neurogenesis.

**Figure 2.**
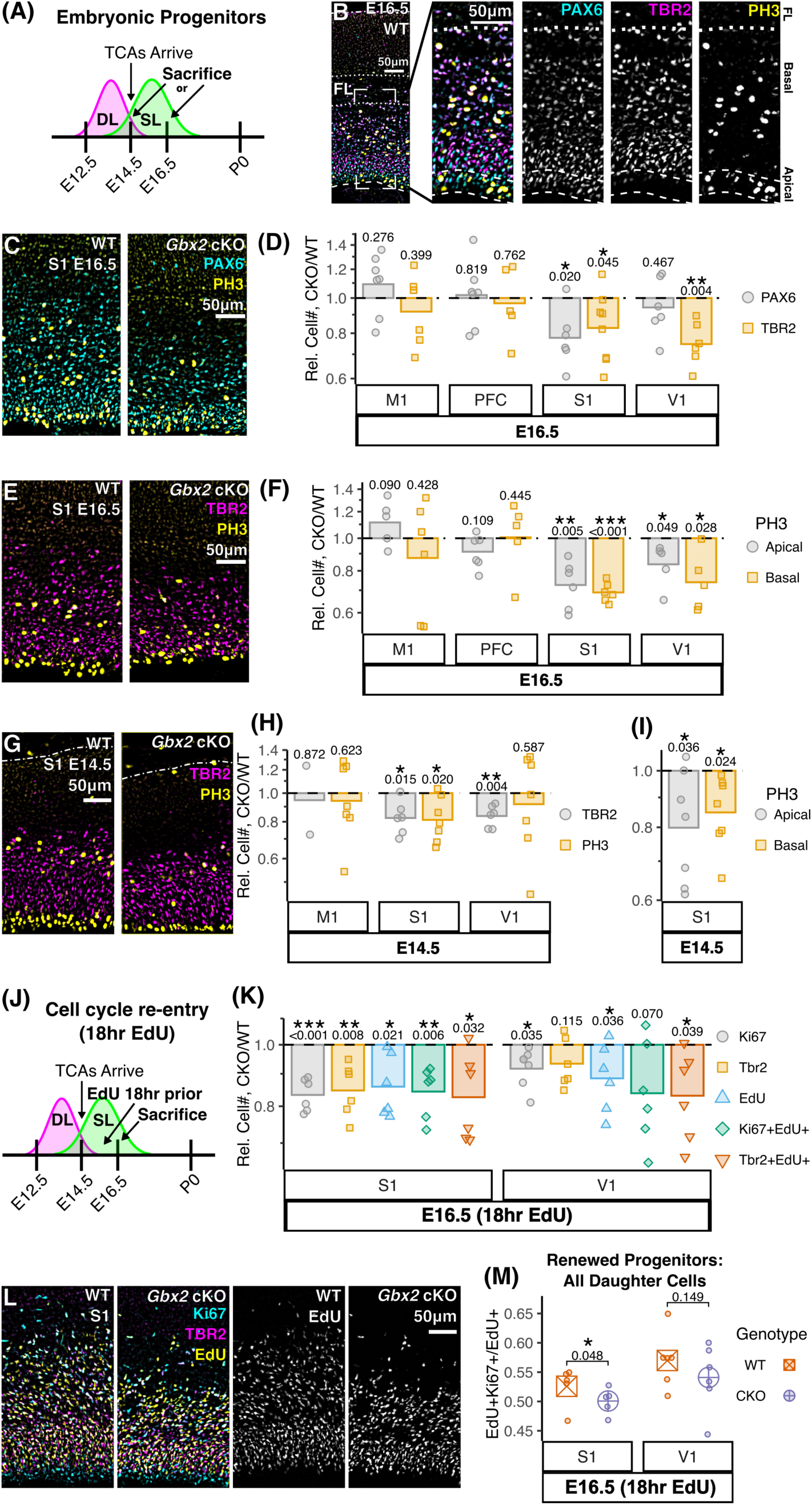
Thalamocortical axons increase progenitor numbers, divisions, and cell cycle re-entry in primary sensory cortex. **(A)** Experimental timeline for B-I. **(B)** S1 wildtype (WT) cortex progenitors—radial glia (PAX6), intermediate progenitors (TBR2), and mitotic cells (PH3)—are below thalamocortical axons (TCAs) in the fiber layer (FL); apical PH3 are strictly radial glia; basal PH3 are PAX6^+^ and/or TBR2^+^. **(C-D)** The number of progenitors is decreased in *Gbx2* cKO mice at E16.5 in S1 (TBR2 & PAX6) and V1 (TBR2), but not in non-sensory cortex M1 (primary motor) or PFC (prefrontal cortex). **(E-F)** At E16.5, both apical and basal progenitors divide (PH3^+^) less in *Gbx2* cKO S1 and V1. **(G-I)** At E14.5 in *Gbx2* cKO mice, TBR2 is decreased in both S1 and V1 and both apical and basal cells divide (PH3^+^) less in S1. **(J)** Experimental timeline for K-M. **(K-L)** E16.5 *Gbx2* cKO mice have decreased numbers of progenitors (Ki67^+^ & TBR2^+^) and fewer cells proceeding through S-phase (EdU^+^) in both S1 and V1. **(M)** More progenitors re-enter the cell cycle (EdU^+^Ki67^+^) in S1 of WT mice compared with *Gbx2* cKO. Matched ratio t-test (D,F,H,I,K) or paired t-test (M) *p*-values shown between WT and *Gbx2* cKO. For (D,F,H,I,K), data shown as relative difference of *Gbx2* cKO to WT littermates (dashed line); points represent individual pairs, bar represents pooled mean. See Fig.S2 for breakdown of multi-channel images.

### Fate decisions among superficial layer neurons is controlled by thalamocortical projections in postnatal sensory cortex

A highly distinct cytoarchitectonic feature of adult primary sensory cortex is the abundance of layer 4 neurons, the main synaptic targets of TCAs derived from corresponding principal sensory nuclei of the thalamus. Synaptic activity of TCAs induces characteristic morphology and patterns of gene expression of layer 4 neurons in juvenile S1 (Narboux-Nême et al., 2012; Li et al., 2013; Pouchelon et al., 2014; De León Reyes et al., 2019) and V1 (Callaway and Borrell, 2011). However, the diversification of layer 4 and layer 2/3 fates is first detected perinatally in immature superficial layer neurons, long before morphological features of each neuronal type become evident (Oishi et al., 2016; Di Bella et al., 2020). It is unknown if this early fate specification is determined strictly by intrinsic gene regulation or is influenced by the thalamic input (Gao et al., 2014; Hou et al., 2019; Llorca et al., 2019; Klingler and Jabaudon, 2020; Llorca and Marín, 2021). Therefore, we next tested if TCAs not only enhance the overall number of superficial layer neurons but also affect the early fate choice between layer 2/3 and layer 4 neurons. For this purpose, we labeled progenitor cells in the S-phase of the cell cycle by EdU at E14.5 or E16.5 (Fig.3A,G), and tracked the fate of neurons that are generated soon after the labeling. We first analyzed each of the cell cohorts at P8 in primary sensory cortex, when excitatory neurons occupy the final laminar positions but may not have fully developed morphological characteristics (De León Reyes et al., 2019).

**Figure 3.**
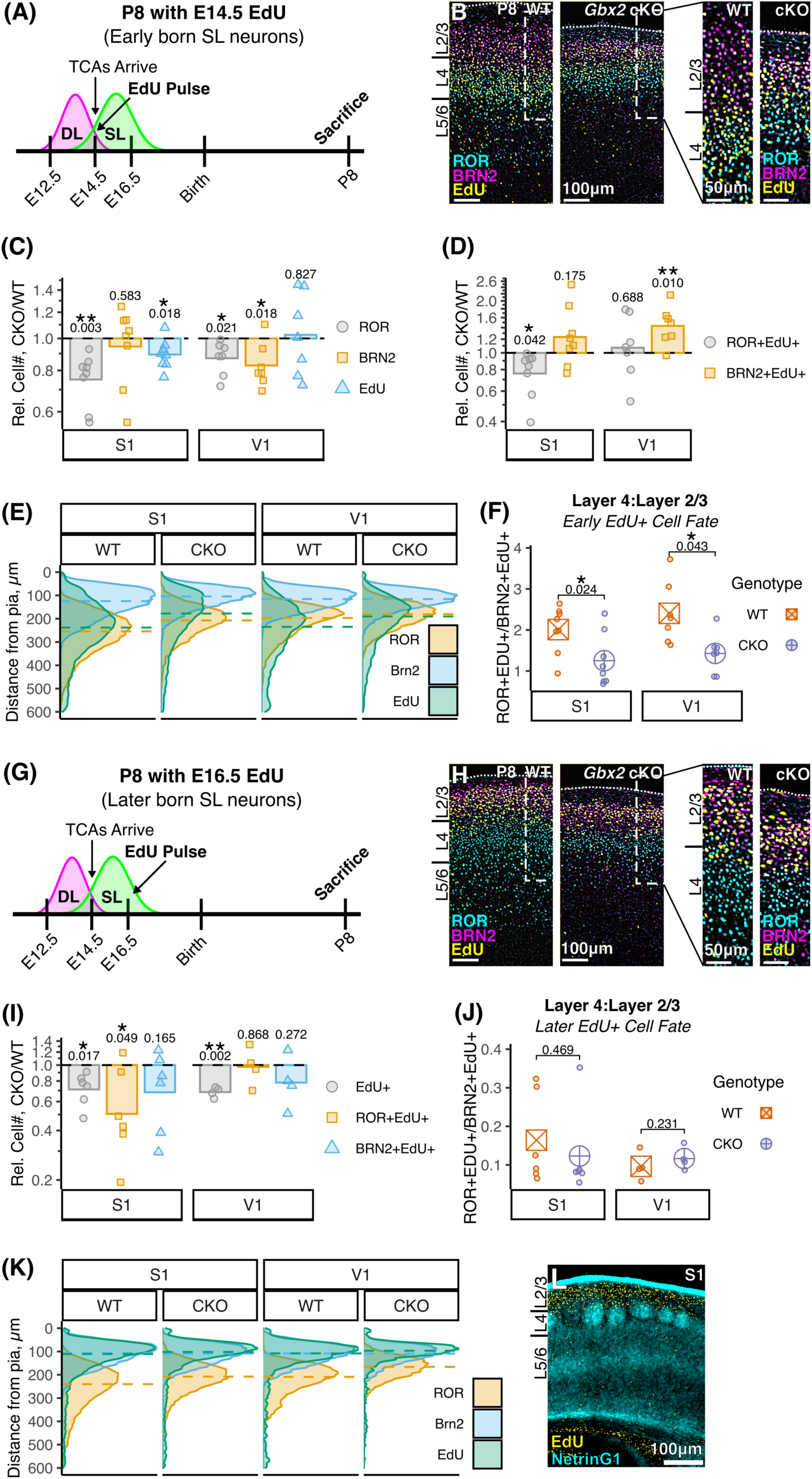
Thalamocortical axons bias early-born superficial layer neurons to become layer 4 instead of layer 2/3. **(A)** Experimental timeline for B-F. **(B)** Distribution of E14.5-labeled EdU+ cells, ROR+ (layer 4) and BRN2+ (layers 2/3) neurons in S1 at P8. The two panels on the right show high magnification views within the dotted lines in the left panels. **(C)** *Gbx2* cKO reduces EdU in both S1 and V1; EdU-injected WT and *Gbx2* cKO have similar changes to the numbers of superficial layer (SL) neurons as in non-injected mice in Fig.1F. **(D)** *Gbx2* cKO reduces ROR+EdU+ cells in S1 and increases Brn2+EdU+ cells in V1. **(E)** EdU+ cells are positioned deeper in S1 and V1 of WT mice compared with *Gbx2* cKO mice. Density curve of distance from pial surface for all cells analyzed in C; dashed line, median distance. **(F)** WT EdU-labeled SL neurons are biased towards ROR and against BRN2 when compared with *Gbx2* cKO. **(G)** Experimental timeline for H-L. **(H)** E16.5-labeled EdU+ cells with SL neuron markers. **(I)** EdU+ cells are reduced in *Gbx2* cKO in both S1 and V1; ROR+EdU+ cells are decreased in S1 *Gbx2* cKO mice. **(J)** E16.5 EdU-labeled neurons are not biased in cell fate by thalamocortical axons (TCAs); note ratio compared to F. **(K-L)** E16.5-labeled EdU cells are positioned superficially, mostly above TCA terminals (NetrinG1) in layer 4. Matched ratio t-test (C,D,I) or paired t-test (F,J) *p*-values shown between WT and *Gbx2* cKO. For (C,D,I), data shown as relative difference of *Gbx2* cKO to WT littermates (dashed line); points represent individual pairs, bar represents pooled mean. See Fig.S3 for breakdown of multi-channel images.

As expected from previous studies (Klingler et al., 2019), the cohort of E14.5-born neurons was mainly found in layer 4 in control S1 (Fig.3B,E,S3A). This cohort of cells was shifted superficially towards the BRN2^+^ layers 2/3 in *Gbx2* cKO mice (Fig.3B,E,S3B), suggesting that postnatal cell position is regulated by the embryonic presence of TCAs. In the cKO mice, the total number of E14.5-EdU^+^ cells was reduced in S1 (Fig.3C), consistent with the reduction of progenitors at E14.5 (Fig.2G-I). The number of E14.5-born neurons that have adopted the layer 4 fate (EdU^+^ROR^+^) was decreased in S1, while E14.5-born layer 2/3 neurons (EdU^+^BRN2^+^) were significantly increased in V1 (Fig.3D). In addition, the ratio of EdU^+^BRN2^+^ cells over the total EdU^+^ cells was increased in both S1 and V1 (Fig.S3C). These changes led to a reduction of the layer 4 to layer 2/3 ratio (EdU^+^ROR^+^/EdU^+^BRN2^+^) in both of these sensory areas of the cKO mice (Fig.3F). Thus, by P8, TCAs normally promote the acquisition of the layer 4 fate at the expense of the layer 2/3 fate in immature neurons generated at E14.5 in primary sensory cortex. This result coincides with the reduced ratio of all layer 4 neurons to layer 2/3 neurons regardless of the birthdate (Fig.1G).

When progenitor cells were labeled by EdU at E16.5 and the brains were analyzed at P8 (Fig.3G), most EdU^+^ cells were located in layers 2/3 in both wildtype and *Gbx2* cKO mice (Fig.3H,K,L, S3D,E). Consistent with the decrease in progenitor cells at E16.5 (Fig.2C-F), the number of EdU^+^ cells was decreased in the cKO mice, and the small number of E16.5-born layer 4 neurons (EdU^+^ROR^+^) was further decreased in S1 (Fig.3I). However, the layer 4 to layer 2/3 ratio was unchanged in the cohort of E16.5-born neurons (Fig.3J). Thus, TCAs increase layer 4 neurons by biasing cell fate in relatively early-born (the E14.5 EdU cohort) superficial layer neurons and by maintaining the production of layer 4 neurons in later-born neurons (the E16.5 EdU cohort).

### Early postmitotic transition of laminar fate is influenced by thalamocortical input prior to direct synapse formation

We next determined whether sensory TCAs affect the laminar fates of E14.5-born neurons earlier than P8, before the formation of the direct synapses between TCAs and prospective layer 4 neurons at around P2 (Agmon et al., 1996; Molnár et al., 2003). In wildtype mice at E16.5, EdU^+^ cells were mainly found in the intermediate zone where IPCs are abundant (Fig.4B,S4B-C); some EdU^+^ cells were found more superficially, in the fiber layer overlapping with the incoming TCAs as well as in the cortical plate. Thus, most E14.5-born neurons are still migrating at E16.5 (Fig.S4A). At this stage, the upper cortical plate (future superficial layer) contained BRN2^+^ neurons, while the lower cortical plate (future deep layer) contained CTIP2, and expression of ROR was mostly contained within the CTIP2^+^ layer (Fig.4C, S4D).

**Figure 4.**
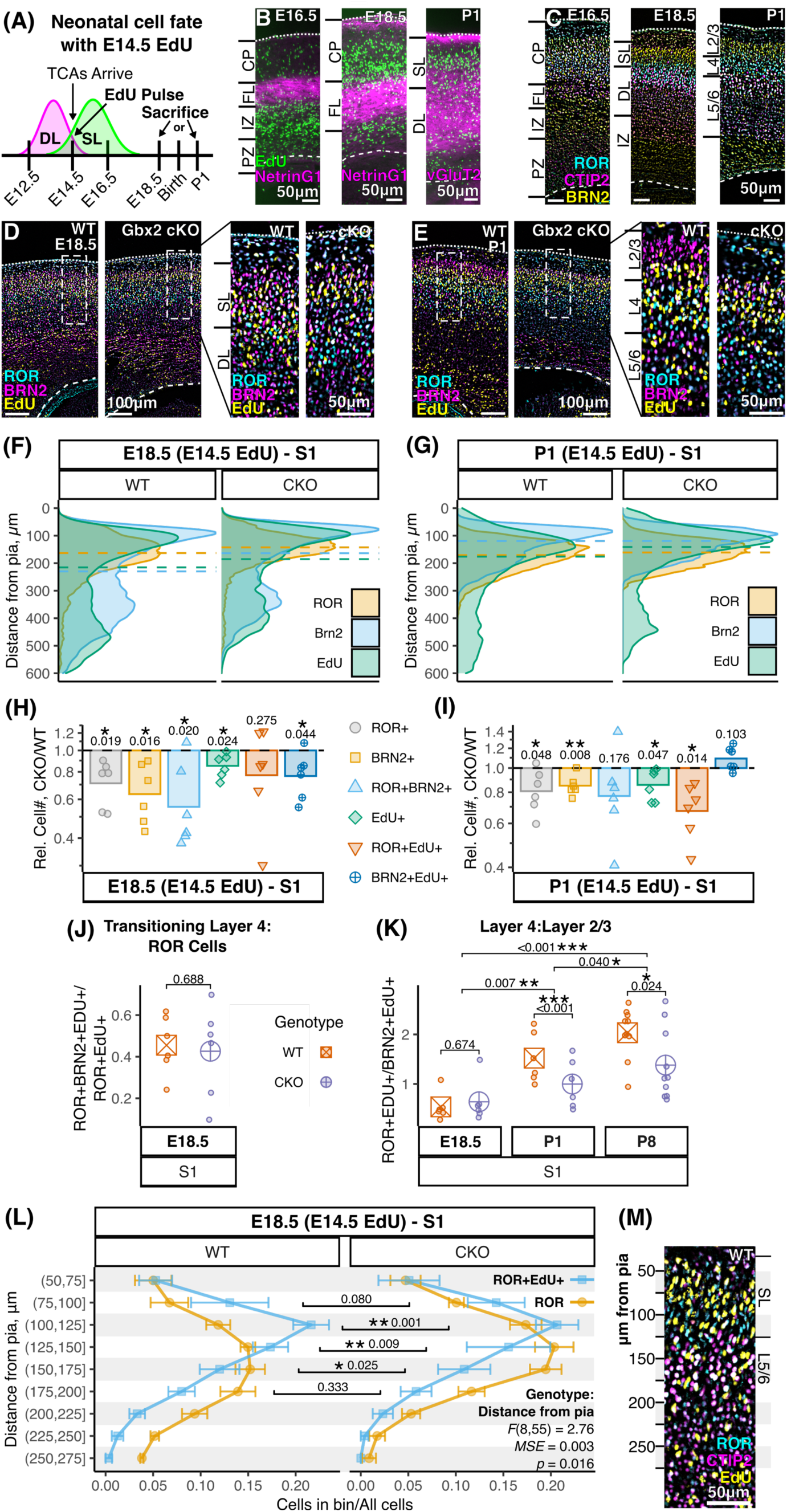
Thalamocortical axons regulate the acquisition of cell fate in the perinatal neocortex. **(A)** Experimental timeline; birth, equivalent to E19.5 or P0. **(B)** Positional relationship between the thalamocortical axons (TCAs) and cells labeled by EdU at E14.5 (see Fig.S4B,E,M for *Gbx2* cKO). At E16.5, labeled cells are mostly below the fiber layer (FL), which contains TCAs (NetrinG1). At E18.5, EdU+ cells are migrating or have migrated through TCAs (NetrinG1). By P1, most labeled cells are in the cortical plate and TCAs (vGluT2) have grown to future layer 4. **(C)** Dynamic changes of ROR expression. At E16.5, ROR is contained in the CTIP2+ layer. At E18.5, ROR still mostly overlaps CTIP2, but a small fraction of ROR+ cells are first found above layer 5. At P1, ROR+ cells form a more distinctive layer between CTIP+ and BRN2+ layers. **(D-E)** Fate analysis of E14.5 EdU-labeled cells analyzed at E18.5 or P1. At E18.5, in both WT and cKO cortex, E14.5-labeled EdU+ cells mostly reside above the layer containing ROR and CTIP2, and express BRN2. At P1, a majority of EdU+ cells are either in the layer expressing ROR (layer 4) or BRN2 (layer 2/3). (**F-G**) Distribution of E14.5 EdU-labeled cells in relation to that of ROR+ and BRN2+ cells at E18.5 or P1. At P1, but not at E18.5, EdU+ cells are positioned more superficially in *Gbx2* cKO mice compared with WT littermates. Density curves of distance from pial surface for all cells analyzed in H-I; dashed line, median distance. **(H)** At E18.5, the number of BRN2+ and ROR+ cells, as well as transitioning ROR+BRN2+ cells, is decreased in *Gbx2* cKO mice; EdU+ and BRN2+EdU+ cells are also decreased. **(I)** At P1, all superficial layer neuron markers (layer 4, ROR; layer 2/3, BRN2), EdU+ cells, and ROR+EdU+ cells are decreased in *Gbx2* cKO mice. **(J)** At E18.5, there is no change in the ratio of E14.5-born transitioning future layer 4 neurons (ROR+BRN2+EdU+) to the total ROR+EdU+ population. **(K)** The ratio of ROR+EdU+ cells over BRN2+EdU+ cells is not significantly changed in *Gbx2* cKO mice at E18.5, but is decreased at P1 and P8. **(L)** There is a significant difference in the percentage of the population (i.e. # cell type in bin / # cell type) between ROR cells and E14.5-born ROR (ROR+EdU+). Linear mixed-effects model for interaction between genotype and distance from pia (repeated measure) shown in plot. *Gbx2* cKO mice have significantly fewer cells just above CTIP2+ layer 5 neurons (C,M,S4I), relevant *p-*values for comparisons between WT and *Gbx2* cKO at a given bin shown in plot. **(M)** ROR+ cells reside both within the deep layers (CTIP2) and above the deep layers; E14.5-born neurons (EdU) that express ROR mostly reside above the CTIP2 layer. CTIP2 layer begins approximately 125-150µm from the pial surface (see Fig.S4I). Matched ratio t-test (H,I) or paired t-test (J,K) *p*-values shown between WT and *Gbx2* cKO. For (H,I), data shown as relative difference of *Gbx2* cKO to WT littermates (dashed line); points represent individual pairs, bar represents pooled mean. See Fig.S4 for breakdown of multi-channel images.

By E18.5, many EdU^+^ cells were found above the fiber layer, mainly within the upper cortical plate overlapping with BRN2 (Fig.4B,D, S4A,E-G). Thus, a majority of E14.5-born neurons migrate past TCAs between E16.5 and E18.5, potentially allowing them to be influenced by these afferent axons without a direct synaptic input from them. At E18.5, a majority of ROR^+^ cells were still within the CTIP2^+^ layer and co-expressed CTIP2 (Fig.4C, S4H,I). The lack of E14.5 EdU labeling in CTIP2^+^ cells (Fig.S4I) suggests that ROR^+^CTIP2^+^ cells are not future layer 4 neurons. At this stage, however, ROR was also expressed in a thin layer of cells above the CTIP2^+^ layer, some of which co-expressed BRN2 (Fig.4C,S4H). This perinatal timing marks the first appearance of layer 4-specific gene expression, which was also detected in single-cell RNA sequencing (Di Bella et al., 2020). In prospective S1 of *Gbx2* cKO embryos at E18.5, EdU^+^ cells were reduced in number compared with wildtype controls, as were the future superficial layer (EdU^+^BRN2^+^) neurons (Fig.4H), reflecting the reduction of progenitor cells at E14.5 (Fig.2G-I). However, compared with P8, a much larger proportion of EdU^+^ cells was positive for BRN2 relative to ROR (compare Fig.S3C and S4J), and within the ROR^+^EdU^+^ population nearly half also expressed BRN2 (Fig.4J), revealing E18.5 as a transitory stage for superficial layer neurons to acquire the laminar fate. In cKO mice there was a significant decrease in ROR^+^EdU^+^ cells closer to the pial surface compared to the bulk of the ROR^+^ cell distribution (Fig.4L-M). Since future layer 5 ROR^+^ neurons were not labeled with EdU (Fig.3B), this shift of ROR^+^EdU^+^ cells away from the pial surface likely represents the reduced specification of future layer 4 (ROR^+^) neurons in the absence of TCAs. Thus, already in the prenatal cortex, TCAs appear to promote the onset of layer 4 characteristics (ROR expression and laminar positioning) in earlier-born superficial layer neurons.

Later at P1, layer 4 and layer 2/3 neurons were separable based on their laminar positions and largely distinct expression of ROR and BRN2, respectively (Fig.4E, S4L), and E14.5-born neurons were superficial to layer 5 (CTIP2) neurons (Fig.S4N). In *Gbx2* cKO S1 cortex, the number of E14.5-born, ROR^+^ cells as well as the layer 4 to layer 2/3 ratio (EdU^+^ROR^+^/EdU^+^BRN2^+^) was decreased compared with wildtype littermates (Fig.4I,K,S4K), indicating a shift of molecular laminar fate from layer 4 towards layer 2/3 without TCAs. In addition, the radial positions of EdU-labeled cells were shifted towards the pia at P1 (Fig.4G, S4M-P). Thus, we conclude that already at the earliest stage when superficial layer neuron fates are distinguishable between layer 4 and layers 2/3, TCAs have promoted the fate of layer 4 neurons at the expense of layers 2/3 neurons among E14.5-born superficial layer neurons.

### VGF is a thalamus-derived molecule that affects progenitor cell division and layer 4 fate in prenatal and perinatal sensory cortex

The numbers of progenitor cells and layer 4 neurons were decreased in prospective primary sensory areas, but not in the motor area or medial prefrontal cortex of *Gbx2* cKO mice. Therefore, we predicted that the responsible TCA-derived molecules that control the number of progenitor cells and laminar fate of immature superficial layer neurons are specifically expressed in principal sensory nuclei of the thalamus, which project axons to primary sensory areas. VGF (non-acronymic) is expressed specifically by nuclei in the thalamus that project to the primary sensory cortex—ventral posterior to S1, dorsal lateral geniculate to V1, and ventral part of the medial geniculate to A1 (Snyder et al., 1998; Bartolomucci et al., 2011; Sato et al., 2012). We found that *Vgf* mRNA started to be expressed in prospective principal sensory thalamic nuclei as early as at E12.5 and continued throughout development (Fig.5B,S5A). VGF and its cleavage products were present at the terminals of TCAs in the neocortex specific to primary sensory areas (Fig.5C,S5B) and application of uncleaved VGF precursor protein enhanced dendritic growth and survival of cortical neurons *in vitro* (Sato et al., 2012). Therefore, we tested if the TCA-derived VGF has a role in progenitor cells and/or immature neurons in the sensory cortex *in vivo* by analyzing thalamus-specific conditional *Vgf* knockout mice (*Vgf* cKO: *Olig3^Cre/+^; Vgf ^flox/flox^*), and compared them with wildtype littermates (*Olig3^+/+^; Vgf ^flox/+^ or Vgf ^flox/flox^*). With the *Olig3^Cre^* allele, *Vgf* was specifically deleted in embryonic thalamus (Fig.S5C).

**Figure 5.**
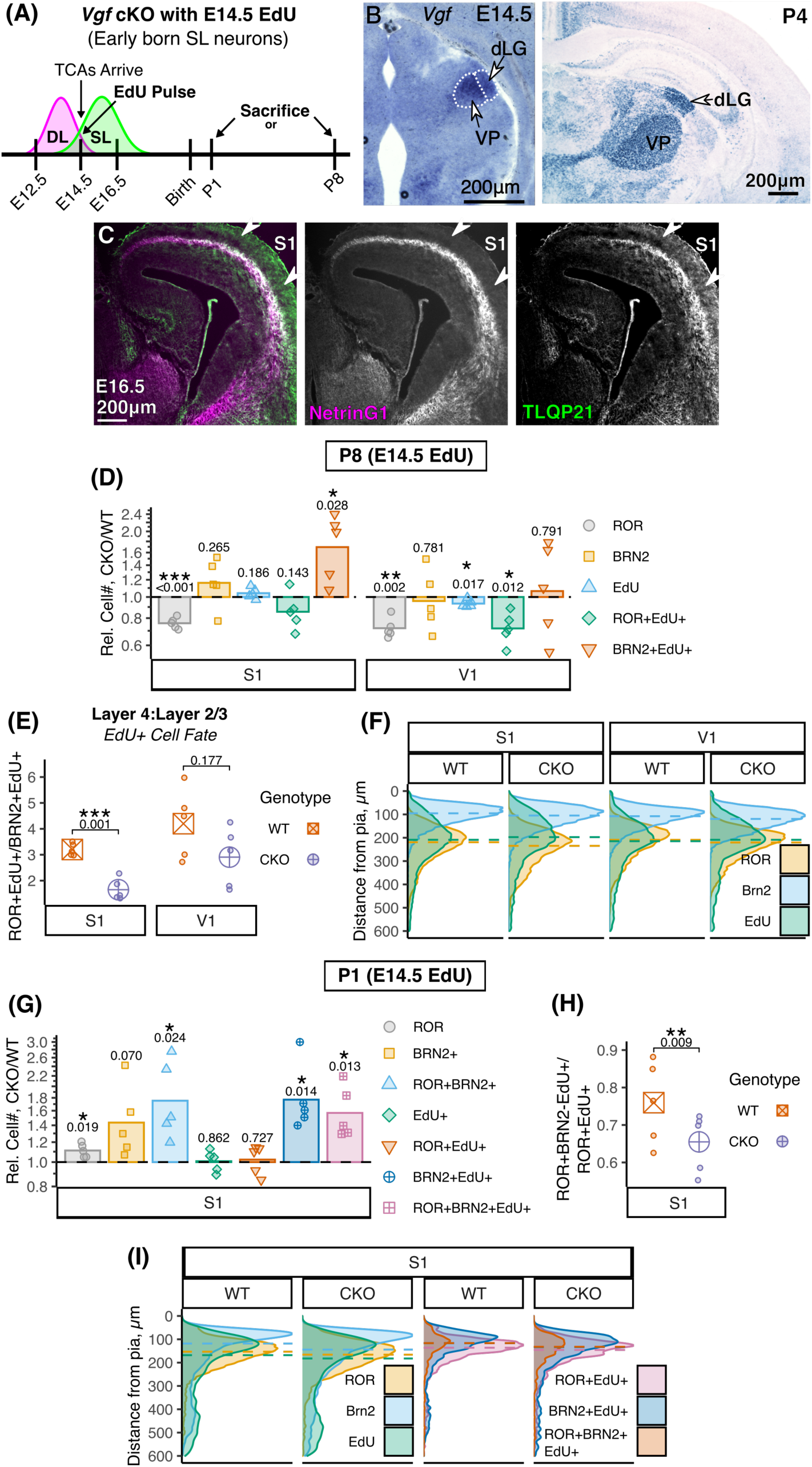
Thalamus-derived, sensory specific VGF peptides increases superficial layer neurons and plays a role in establishing separate ROR+ and BRN2+ cell populations. **(A)** Experimental timeline. (B) *In situ* hybridization for the *Vgf* mRNA. Principal sensory thalamic nuclei (VP, ventral posterior; dLG, dorsal lateral geniculate) express *Vgf*. High expression of *Vgf* remains isolated to sensory nuclei in the postnatal brain. **(C)** At E16.5, TCAs (NetrinG1) contain high levels of a VGF peptide, TLQP21, in future sensory areas (S1). **(D)** At P8, *Vgf* cKO mice injected with EdU at E14.5 have reduced numbers of layer 4 ROR+ neurons in both S1 and V1 compared with WT mice. EdU is slightly decreased in V1 of *Vgf* cKO mice. *Vgf* cKO mice have more Brn2+EdU+ cells in S1 and fewer ROR+EdU+ cells in V1. **(E)** VGF biases E14.5-born neurons to become layer 4 (ROR+) compared to layers 2/3 (BRN2+) in S1 with a trend towards this ratio in V1. **(F)** Radial distribution of E14.5 EdU+ cells is not different between WT and *Vgf* cKO mice at P8. Density curve of distance from pial surface for all cells analyzed in D; dashed line, median distance. **(G)** At P1, *Vgf* cKO mice have increased ROR+, BRN2+, and ROR+BRN2+ cells as well as more BRN2+EdU+ and ROR+BRN2+EdU+ cells after EdU injection at E14.5. **(H)** At P1, *Vgf* cKO mice have fewer E14.5-born (EdU+) ROR+ neurons that transition to fully specified layer 4 (ROR+BRN2-) neurons. **(I)** At P1, EdU, ROR, and BRN2 distribution is not different between WT and *Vgf* cKO mice. More ROR+BRN2+EdU+ cells are present in *Vgf* cKO mice and the increased overlap of ROR+EdU+ and BRN2+EdU+ cells is likely to account for this. Density curve of distance from pial surface for all cells analyzed in G; dashed line, median distance. Matched ratio t-test (D,G) or paired t-test (E,H) *p*-values shown between WT and *Gbx2* cKO. For (D,G), data shown as relative difference of *Gbx2* cKO to WT littermates (dashed line); points represent individual pairs, bar represents pooled mean. See Fig.S5 for breakdown of multi-channel images.

At P8, *in situ* hybridization for *RORβ* mRNA showed thinning of layer 4 in the barrel field of S1 (Fig.S5D). With immunohistochemistry combined with EdU injection at E14.5, the numbers of ROR^+^ layer 4 neurons in S1 and V1 were reduced in P8 *Vgf* cKO mice (Fig.5D), and the number E14.5-born layer 4 neurons (EdU^+^ROR^+^) was also reduced in V1. However, unlike in *Gbx2* cKO, the total number of E14.5 EdU-labeled cells was unchanged (Fig.5D), suggesting that for the cohort of E14.5-born neurons, reduction of ROR^+^ cells was not caused by the reduced overall production of superficial layer neurons. At E16.5, the numbers of both radial glia and intermediate progenitor cells were unchanged in *Vgf* cKO mice, although the number of basally dividing cells (PH3+) cells was reduced in both prospective S1 and V1 (Fig.S5E,F), which indicates that VGF might control the cell cycle length of intermediate progenitor cells in sensory cortex. Thus, the lack of TCA-derived VGF accounts for only part of the phenotype seen in TCA-ablated (*Gbx2* cKO) cortex. At P8, the EdU^+^ROR^+^/EdU^+^BRN2^+^ ratio was decreased in *Vgf* cKO S1 (Fig.5E) even though, unlike *Gbx2* cKO, E14.5-born cells were not distributed differently among layers 4 (ROR) and 2/3 (BRN2) (Fig.5F), suggesting that the TCA-derived VGF promotes the layer 4 fate at the expense of the layer 2/3 fate regardless of the overall cell number or position. We therefore investigated if VGF, like the TCAs themselves, already affects the fates of superficial layer neurons by neonatal stages. Like at P8, the number of neurons labeled by E14.5 EdU injection was not changed at P1 (Fig.5G), nor was the radial distribution of EdU^+^ cells different compared with wildtype littermates (Fig.5I). However, *Vgf* cKO mice had an increased number of E14.5-born neurons that express BRN2 as well as those that express both BRN2 and ROR (Fig.5G). Considering that prospective layer 4 neurons appear to transition from BRN2^+^ROR^-^ through BRN2^+^ROR^+^ to BRN2^-^ROR^+^ during perinatal stages (Fig.S4A), the observed changes in P1 *Vgf* cKO S1 cortex suggest that VGF is required for the timely transition from the transitory, BRN2^+^ROR^+^ state to the fully specified layer 4 (BRN2^-^ROR^+^) state likely by suppressing the expression of BRN2. Consistent with this conclusion, we found a decrease in the EdU^+^BRN2^-^ ROR^+^/EdU^+^ROR^+^ ratio in the cKO brains (Fig.5H).

## Discussion

The current study has revealed two distinct roles of TCAs in the early stages of the formation of cytoarchitecture in the primary sensory cortex. First, TCAs enhance neurogenesis during embryonic stage by increasing the number of progenitor cells that produce superficial layer neurons. In addition, TCAs promote the specification of the layer 4 fate in immature superficial layer neurons at the expense of the layer 2/3 fate by the neonatal stage. These two early roles are consistent with the physical proximity of TCAs to both progenitor cells and immature neurons at the relevant developmental timing before direct synapses are formed between TCAs and layer 4 neurons. Furthermore, we found that the thalamus-derived molecule VGF partially contributes to those roles, implying that multiple molecules derived from thalamic afferents play a role in neurogenesis and fate specification in early sensory cortex.

Afferent control of neurogenesis in the target region of the developing central nervous system has long been known in the invertebrate visual system, in which retinal axons control cell cycle progression and subsequent differentiation of neurons in their target brain region via the secreted protein Hedgehog (Selleck et al., 1992; Huang and Kunes, 1996). In zebrafish, descending projections of diencephalic dopaminergic neurons enhance the production of motor neurons in the spinal cord (Reimer et al., 2013). However, little is known about the direct regulation of neurogenesis by afferent projections in mammalian brains. Protracted neurogenesis in the mammalian neocortex and early projection of TCAs allow close interactions between the afferents and progenitor cells in embryonic cortex (Reillo et al., 2017). Despite the evidence from eye removal studies in monkeys and ferrets (Rakic, 1988; Dehay et al., 1989; Rakic et al., 1991; Dehay et al., 1996; Reillo et al., 2011), direct demonstration of the roles of the TCAs on progenitor cells was lacking. Our current study utilized a mouse model in which few TCAs reach the cortex and found that both radial glia and intermediate progenitor cells are reduced in number. In addition, progenitor cells are more likely exit the cell cycle in the absence of normal TCA input in prospective primary sensory areas in embryonic cortex. Further, we found reduced mitosis of neural progenitor cells in late stages of neurogenesis in mice lacking the TCA-derived protein VGF and its processed neuropeptides. However, lack of VGF did not cause the same extent of the defects in neurogenesis observed in mice lacking TCAs, indicating that other thalamus-derived molecules also control the division of cortical progenitor cells.

In the current study, thalamic afferents were found to promote the early specification of layer 4 fate in the primary sensory cortex. Previous studies indicated that RORβ and BRN2 have mutually repressive interactions that result in the emergence of gene expression profile and morphological characteristics of layer 4 versus layer 2/3 neurons during perinatal development (Oishi et al., 2016; Clark et al., 2020), and that an upstream intrinsic transcriptional network regulates the expression of RORβ and BRN2 (Hou et al., 2019). However, it was unknown if this early fate specification of immature superficial layer neurons is determined entirely by intrinsic mechanisms or is influenced by the incoming thalamic input in sensory cortex. Our study demonstrates a role of TCAs in this process and the involvement of the thalamus-derived molecule VGF. Our results do not exclude the possibility that TCAs modify the identity of radial glia or intermediate progenitor cells in prospective sensory areas so that they have a longer time window and/or a higher likelihood of generating layer 4 neurons as opposed to layer 2/3 neurons. However, a recent single-cell RNA sequencing study showed that newborn superficial layer neurons lack distinct gene expression patterns that separate layer 4 and layer 2/3 fates until the perinatal stage, suggesting that the diversification and identity acquisition of neuronal subpopulations occur postmitotically (Di Bella et al., 2020). This is consistent with previous studies (Oishi et al., 2016; De León Reyes et al., 2019). Therefore, it is more likely that TCAs bias immature postmitotic neurons to take on the layer 4 fate within primary sensory cortex. Interestingly, most neurons labeled by EdU injection at E16.5 continued to express BRN2 and migrated to form layers 2/3 in S1 even with TCAs in their vicinity throughout their early development. If TCAs act on postmitotic neurons to influence their fate, how they preferentially bias E14.5-born but not E16.5-born, neurons to take on the layer 4 fate remains to be elucidated.

Our analysis of conditional *Vgf* knockout mice provides evidence that the thalamus-derived molecule VGF mediates distinct roles of the thalamic axons on neocortical development. A recent unpublished study (Sato et al., 2021) showing that germline deletion of *Vgf* reduces the number of ROR-expressing cells in S1 and V1 at P7 is consistent with our results on thalamus-specific *Vgf* deletion. The VGF precursor protein is cleaved into at least a dozen biologically active peptides (Bartolomucci et al., 2011), and little is known about the roles of individual VGF peptides in neural development. One candidate is the C-terminal VGF peptide TLQP-62, which controls hippocampal neurogenesis (Thakker-Varia et al., 2014) as well as depression (Jiang et al., 2018) and nerve injury-induced hypersensitivity (Skorput et al., 2018), although the identity of its receptor is currently unknown. Another C-terminal peptide, TLQP-21 (Bartolomucci et al., 2006; Riedl et al., 2009), is localized in TCAs (Fig.5C) and modulates microglial function through its receptors C3aR1(Hannedouche et al., 2013; Cero et al., 2014; Doolen et al., 2017; El Gaamouch et al., 2020) and C1qBP (Elmadany et al., 2020). Thus, these peptides might have direct or indirect effects on newborn neurons or progenitor cells during embryogenesis and early postnatal cortical development.

In summary, our present study provides a mechanism by which afferent input from the thalamus complements the intrinsic program in early cortical development, allowing the sequential generation of diverse neuronal types in an area-specific manner. Further studies are expected to reveal the spatially and temporally regulated molecular interplay between the thalamic input and cortical cells across different areas.

## Acknowledgments

We thank Earl Parker Scott, Thomas Bao, Emma Breyak, Ellie Wheeler and Xiangyu Zou for technical assistance; Matt Simon, Lynn Wang, Alicia Cowles, Troy Liu, Yuria Shimizu, Samantha Dabruzzi and Peter Goncharov for contribution to early phase of the study; Marija Cvetanovic, Zhe Chen, Linda McLoon and Steven McLoon for critical reading of the manuscript; Lucy Vulchanova for anti-TLQP-21 antibody; Henk Stunnenberg for anti-RORβ antibody.

## Author contributions

TM and YN designed the experiments; TM, JR, JS and YN performed the experiments; TM analyzed the data with the help of JR and JS; SS provided *Vgf^flox^* mice and edited the manuscript. TM and YN wrote and edited the manuscript.

## Supplementary Materials

### Materials and Methods

#### Mice

Care and experimentation on mice were done in accordance with the Institutional Animal Care and Use Committee at the University of Minnesota. Noon of the day on which the vaginal plug was found was counted as embryonic day 0.5 (E0.5), and the day of birth (E19.5) was designated as postnatal day 0 (P0). To generate mice with a conditional deletion of thalamocortical axons (TCAs), *Olig3^Cre/+^*;*Gbx2^null/+^* mice were bred with *Gbx2^flox/flox^* mice as in (Vue et al., 2013). *Gbx2^flox/flox^* mice (Li et al., 2002) were obtained from Jackson Laboratory. *Olig3*^+/+^; *Gbx2^flox/+^* were used as wildtype (WT) littermates and compared to *Olig3^Cre/+^*; *Gbx2^null/flox^* as *Gbx2* conditional knockout mice (cKO). Mice were kept in C57BL/6 background and embryos or pups of either sex were used. Mice are matched within littermates of WT and cKO to control for small differences between litters; therefore, litters without both a cKO and a WT pup were not analyzed.

As with *Gbx2* cKO mice, we generated mice with a conditional deletion of *Vgf* from TCAs. *Olig3^Cre/+^*;*Vgf ^flox/+^* mice were bred with *Vgf ^flox/flox^* or *Vgf ^flox/+^* mice (*66*). *Vgf ^flox^* mice. *Olig3^+/+^; Vgf ^flox/(flox or +)^* or *Olig3^Cre/+^; Vgf ^+/+^* were used as WT littermates and compared to *Olig3^Cre/+^*; *Vgf ^flox/flox^* as *Vgf* cKO mice. As mice *Gbx2* cKO mice, *Vgf* cKO mice were kept in C57BL/6 background, pups of either sex used, and mice were matched littermates for comparison between cKO and WT.

#### EdU incorporation

For 5-ethynyl-2’-deoxyuridine (EdU, Carbosynth, Compton UK) incorporation experiments, mice were administered 50μg EdU/g body weight dissolved in saline via intraperitoneal injection. Pregnant female dams were injected with EdU at the desired stage between 10am and 1pm, and sacrificed 18-hours post injection on E16.5 for 18-hour EdU incorporation experiments. For cell fate analysis, mice were injected with EdU on E14.5 or E16.5 and sacrificed at E16.5, E18.5, P1 or P8 as needed.

#### Tissue preparation

Embryonic day 14.5 (E14.5) mice were removed from the dam and heads were immersion-fixed in 4% paraformaldehyde (PFA) (dissolved in 0.1M phosphate buffer (PB)) for 30 minutes. E16.5 mice were removed from the dam and intracardiacally perfused with 4% PFA, brains dissected in 0.1M PB, and immersion fixed in 4% PFA for 30 minutes. Postnatal mice were intracardiacally perfused by 4% PFA, brains dissected in 0.1M PB, and immersion fixed in 4% PFA for 45 minutes or 60 minutes for P1 and P8, respectively. P8 brains were first perfused by 0.1M PB before PFA. All brains were then submerged in 30% sucrose/ 0.1M PB overnight at 4°C, and frozen in TissueTek OCT compound on dry ice. Frozen brains were stored at −80°C until usage.

#### *In situ* hybridization

*In situ* hybridization using digoxigenin-labeled probes was performed based on (Vue et al., 2007). Embryonic brains were cut with a cryostat at 20μm thickness and P8 brain at 40μm thickness. Sections on slides were left to dry on a slide warmer for 30 minutes prior to starting pre-treatment of sections.

#### Immunohistochemical staining and EdU detection

Immunohistochemical staining was performed based on (Vue et al., 2007). Embryonic and postnatal day 1 brains were cut with a cryostat at 20μm thickness and P8 brain at 40μm thickness. Sections on slides were left to dry on a slide warmer for 30 minutes prior to starting immunohistochemistry. Matched WT and cKO littermates were sectioned onto the same slides to reduce variability of immunohistochemistry between slides. Sections were then rinsed in 0.1M phosphate-buffered saline (PBS), immersed in boiled 10mM citrate buffer (pH 6) for 5 minutes prior to blocking with 3% donkey serum / 0.3% Triton X-100/ PBS for 1 hour. Primary antibodies were then added at appropriate dilutions and incubation was performed overnight at 4°C. The following primary antibodies were used:

| *Protein* | *Dilution* | *Manufacturer* | *Product #* | *Species* |
| --- | --- | --- | --- | --- |
| PH3 | 100 | Millipore | 05-806 | Mouse |
| Ki67 | 100 | BD Biosciences | 550609 | Mouse |
| PAX6 | 100 | BioLegend | 901302 | Rabbit |
| PAX6 | 100 | BD Biosciences | 561462 | Mouse |
| TBR2 | 200 | Abcam | ab23345 | Rabbit |
| vGLUT2 | 100 | Synaptic Systems | 135403 | Rabbit |
| NetrinG1 | 100 | R&D Systems | AF1166 | Goat |
| CTIP2 | 200 | Abcam | Ab30455 | Rat |
| ROR | 100 (50 w/ EdU) | R&D Systems | PP-H3925-00 | Mouse |
| BRN2 | 100 | Santa Cruz | sc-6029 | Goat |
| CC3 | 100 | Cell Signaling | D175 | Rabbit |
| TLQP21 | 200 | Vulchanova lab | Riedl et al. 2009 | Rabbit |
| VGF(R15) | 100 | Santa Cruz | sc-10383 | Goat |

On the second day, after washing in PBS, sections were incubated with fluorochrome-conjugated secondary antibodies (Cy2, Cy3, or Cy5, from Jackson ImmunoResearch, West Grove, PA) for 1 hour, followed by DAPI counterstaining. After dehydration in ascending concentrations of ethanol and clearing in xylene, slides were mounted in DPX mounting medium (EM Sciences, Hatfield, PA).

If EdU was to be visualized then slides were treated under the aforementioned immunohistochemical protocol and then after the treatment of secondary antibodies slides were washed with PBS three times. Then, a solution of 100mM Tris pH7.5/ 4mM CuSO_4_ / 100mM sodium ascorbate (freshly prepared)/ 5μM sulfo-cy5 azide (Lumiprobe, Hunt Valley, MD) was applied to slides for 30 minutes protected from light. Slides were rinsed twice with PBS and then treated with DAPI stain and dehydrated as with immunohistochemistry protocol above.

#### Imaging

For cell counting, images of sections that underwent immunohistochemistry were taken using an E800 upright microscope (Nikon, Tokyo, Japan) with a Retiga EXi camera (Qimaging, Burnaby, BC, Canada). Early images were taken using Open Lab software (Improvision, Lexington, MA) in 12-bit grayscale TIFF format. More recently, we used Micromanager (Edelstein et al., 2014). Micromanager images were obtained using the Multi-Dimensional Acquisition tool and saved as 12-bit OME-TIFF files. The exposure time for a given antibody was selected to achieve the maximum signal-to-noise ratio without reaching maximal intensity; this exposure time was identical for each slide with the same antibody within a matched pair. Images were taken in the order of Cy3, Cy2, Cy5 to avoid cross-channel excitation.

#### Binning

Images are entirely processed within the FIJI (Fiji is just ImageJ) distribution of ImageJ (Schindelin et al., 2012). With Open-Lab Images, individual TIFF files for each channel were opened and combined into a composite stack (see code/software: FIJI_BinningMacro_OpenLab.ijm). Then, a rectangular ROI (region of interest) was overlaid onto the image from pial surface to ventricle and at a width of either 100µm (embryonic non-sensory cortex), 200µm (mid-embryonic sensory cortex), and 400µm (late-embryonic and postnatal cortex). Images acquired with micro-manager were opened and the bin overlayed (FIJI_BinningMacro_MicroManager.ijm). An embryonic and early postnatal mouse atlas (Paxinos et al., 2006) was used to select the bin position to fit within the putative cortical region of interest based on anatomical landmarks. In addition, immunohistochemical markers, such as TBR2 during embryonic development, were used to also orient location of the bin.

#### Thresholding

Images were processed with a custom ImageJ macro, which would batch-process all images with identical parameters across paired littermates. Originally, the process was 1) simple paraboloid background subtraction 2) normalization of signal intensity 3) thresholding by manually determined percentage 4) default watershed algorithm (FIJI_Thresholding-ITCN.ijm). Over time, the complexity of images (overlapping nuclei, multi-channel colocalization, poor signal-to noise) increased which meant there was continual development of the code (FIJI_Thresholding_WatershedSegmentation.ijm). The final macro can be applied to all images and advantageously preserves near maximum width of cell nuclei signal while also having efficient separation of unique nuclei; previous analysis using the original method were cross validated with the new method to ensure reproducibility of the dataset. First, a convoluted background subtraction was applied with a median filter radius of approximately the diameter of a larger cell of interest, in pixels (Biovoxxel Toolbox: Convoluted Background Subtraction plug-in). The image was then filtered with a median blur to reduce image sensor and/or electrical noise produced by the camera but to preserve edges. A prominence map (peak-valley distance mapping) was used to produce a Voronoi segmented particle image from prominence peaks. The original image was then converted to 32-bit floating point values and divided by the maximal intensity within the region of interest to normalize the image; this was meant to reduce signal variability incurred by immunohistochemistry between sections and slides as well as to avoid normalizing against auto-fluorescent signal such as from dust or cross-reactivity with non-parenchymal tissue such as the choroid plexus. An Otsu threshold algorithm was run on the image using a value which created maximal separation between cells without losing cells of an experimenter-determined minimal intensity; the lower threshold value was usually between 10 and 25% of the maximum normalized intensity. Thresholding was used to remove variability and bias for manual selection of cell nuclei by an individual. Finally, a prominence-based algorithm for watershed was employed by calculating the logical AND operation of the Voronoi segmented prominence map and the thresholded image; the resulting output is more accurate in separating cells compared to the original method, especially because particle size and overlapping nuclei directly reduce efficacy of traditional watershed algorithms. Finally, an optional removal of small contiguous pixels below an area of five pixels was used for antibodies with weaker signal-to-noise (Morphology: Particles8 plug-in).

#### Colocalization

To determine double and triple positive cells, an optional section within the thresholding batch macro or a separate batch macro (FIJI_Coloc_Subtractor.ijm) was utilized. To identify cells double-positive for two different antibodies, one binary channel of the image stack was identified as the “Object” and a different binary channel was the “Selector”. The Binary Feature Extractor plug-in was used to set a percent overlap for the Object on top of the Selector; nuclei of the Selector were reconstructed on a new image channel if the criteria for overlap was met. Using the Analyze Particles tool, the outlines of each nuclei ROI was compared against the original images to confirm that the overlap percentage was not reconstructing aberrant cell nuclei; the overlap percentage was set to between 35% and 70% with intent to be strict on reconstructing cells and thus not all experimenter-observed overlapping cells were reconstructed at the cost of ensuring reconstructed cells were correct. Triple-positive cell nuclei were identified by using a double-positive reconstructed image as the Selector and the third channel as the Object. In addition, the separate colocalization code can also be used to identify cells which are only single positive by subtracting double positive reconstructed cells from the original binary cell image (FIJI_Coloc_Subtractor.ijm).

#### Cell counting

##### With ImageJ/FIJI

Original cell counts were done with the ITCN plug-in for ImageJ which consists of most non-EdU postnatal experiments. Binary and binned images were randomly renamed to remove all identifying information and blind experimenters to genotype. A radius was selected to ensure only single counting of cells. ITCN requires manual selection of the area.

##### With R

As cell counting difficulty increased, such as with more channels and colocalized images, and thresholding sensitivity and accuracy increased, a more autonomous method of cell counting was desired. With improved thresholding, a contiguous set of binary “1” pixels corresponded accurately to a single cell (see Thresholding section). Therefore, setting a unique identifier to a contiguous pixel unit (e.g. from 1 – n) allows for automated cell counting with code (R_Image_Analysis.Rmd). All binary thresholded images were imported into R and represented as a binary matrix. Image metadata is extracted from a consistent file name formula. Contiguous pixel units (a cell) are labeled (ImageR package) with a unique identifier and features such as pixel position, center of mass, unit area, and relative position of the cell.

The data is collected for each image within the imported stack and organized into different tables; these tables are exported as .csv files available for manual inspection or statistical analysis with R.

##### Cell distribution

Binary images were straightened with respect to the ventricular and pial surface of the neocortex if necessary (FIJI_Straighten.ijm) so that either the x- or y-axis could be used for a measurement of the distance of a cell nuclei from the cell axis. Using R (R_Image_Analysis.Rmd), .tiff images were loaded into the environment and represented as a binary matrix. Images were preprocessed in FIJI to set the bounds of the maxima and minima to the pial and ventricular surfaces. Individual nuclei were determined by contiguous pixel sets using a labeling function. Then, average x- and y-coordinates were determined for the location of each nucleus relative to an axis. Finally, distance was normalized to the maximal and minimal distance along the ventricle-to-pia axis and saved as relative distance values. Datasets were saved as .csv files and compared statistically using R.

#### Experimental Design and Statistical Analyses

Littermates were matched for experiments and analysis between *Gbx2* cKO or *Vgf* cKO and WT mice to control for immunohistochemical variability between pairs; WT mice are the control. Both males and females were used in this study and preliminary data showed no differences between males and females, so male and female mice were not separated for analysis. At least five matched pairs were used for each analysis to achieve a statistical power > 0.8. For each neocortical area of interest, multiple brain sections (minimum of three) were analyzed and imaged; data from the same neocortical area within one mouse was averaged to have a representative cell count for the area using R. Ratios of cell types, such as the ratio of cells that are A^+^B^+^/A^+^, were determined from the averaged representative cell counts and not from a per-section basis.

#### Statistics

Matched ratio t-test was used for comparing the logarithm of the ratiometric average representative cell counts between *Gbx2* cKO or *Vgf* cKO and WT littermates. A paired t-test was used to compare ratios, such as A^+^B^+^/A^+^, between *Gbx2* cKO or *Vgf* cKO and WT littermates. For Figure 4K, a mixed linear model was used to compare the ROR+EdU+/BRN2+EdU+ between *Gbx2* cKO mice and WT mice, with E18.5, P1, and P8 as a within subject factor. For distribution analysis, multiple sections of the same neocortical area within one mouse were pooled to increase accuracy of cell distribution. A Kolmogorov-Smirnov test was used to test for significant differences in the distributions of a cell type between *Gbx2* cKO and WT mice, but data is not shown. For Figure 3L, a linear mixed-effects model was used to compare the difference in the percentage of the population (i.e. # cell type in bin / # cell type) to test for interaction between genotype and distance from pia as a repeated measure. A *p-* value < 0.05 on a two-tailed analysis determined a significant difference between groups for all statistical tests. R_Distribution_Statistics.Rmd and R_CellCount_Statistics-ITCNcompatible.Rmd was used for statistical tests and creation of plots for cells analyzed using R or FIJI/ITCN, respectively.

#### Code/Software

##### FIJI

The FIJI package based on ImageJ2 was utilized for all processing of image data directly from the microscope (Schindelin et al., 2012; Rueden et al., 2017). Custom FIJI macros were written using the ImageJ Macro scripting language and are available on a GitHub repository (link). The ITCN plugin was used for cell counting binary images (Kuo and Byun, https://imagej.nih.gov/ij/plugins/itcn.html). The BioVoxxel Toolbox was used for ‘Convoluted Background Subtraction’ for improved background subtraction and ‘Binary Feature Extractor’ for colocalization analysis (Brocher, http://www.biovoxxel.de/development/). The Morphology plugins were used for ‘Particles8’ to remove small binary features not meeting pre-determined pixel sizes (Landini, http://www.mecourse.com/landinig/software/software.html). All code created for this project are freely available for use on GitHub: (https://github.com/TimMonko/NeurogenesisThalamocorticalPaper/) A table of currently used code is found below, but defunct code is also referenced in methods and available.

|  |  |
| --- | --- |
| FIJI_BinningMacro_Micromanager.ijm | Combine multi-channel images and bin |
| FIJI_Thresholding_WatershedSegmentation.ijm | Thresholding and optional colocalization |
| FIJI_Straighten.ijm | Straighten images for distribution analysis |
| FIJI_Coloc_Subtractor.ijm | Identify overlapping and non-overlapping nuclei |
| R_Image_Analysis.Rmd | Distribution and counting of cells |
| R_Distribution_Statistics | Statistical comparisons distribution and cell counts |

*R.* The R Project (R Core Team https://www.r-project.org/) was utilized for mathematical operations on binary images processed in FIJI and for statistical analysis of both FIJI and R-derived datasets (Team, 2013) (available at https://www.r-project.org/)). RStudio (RStudio Team http://www.rstudio.com/) was used as the integrated development environment. In addition, the following packages were used: tidyverse, magick, imageR, broom, magick, extrafont, ggh4x, svglite and rstatix. All graphs were made in R utilizing Tidyverse’s ggplot2 and were made publishable with the svglite and broom packages.

## Supplementary Figures

**Figure S1.**
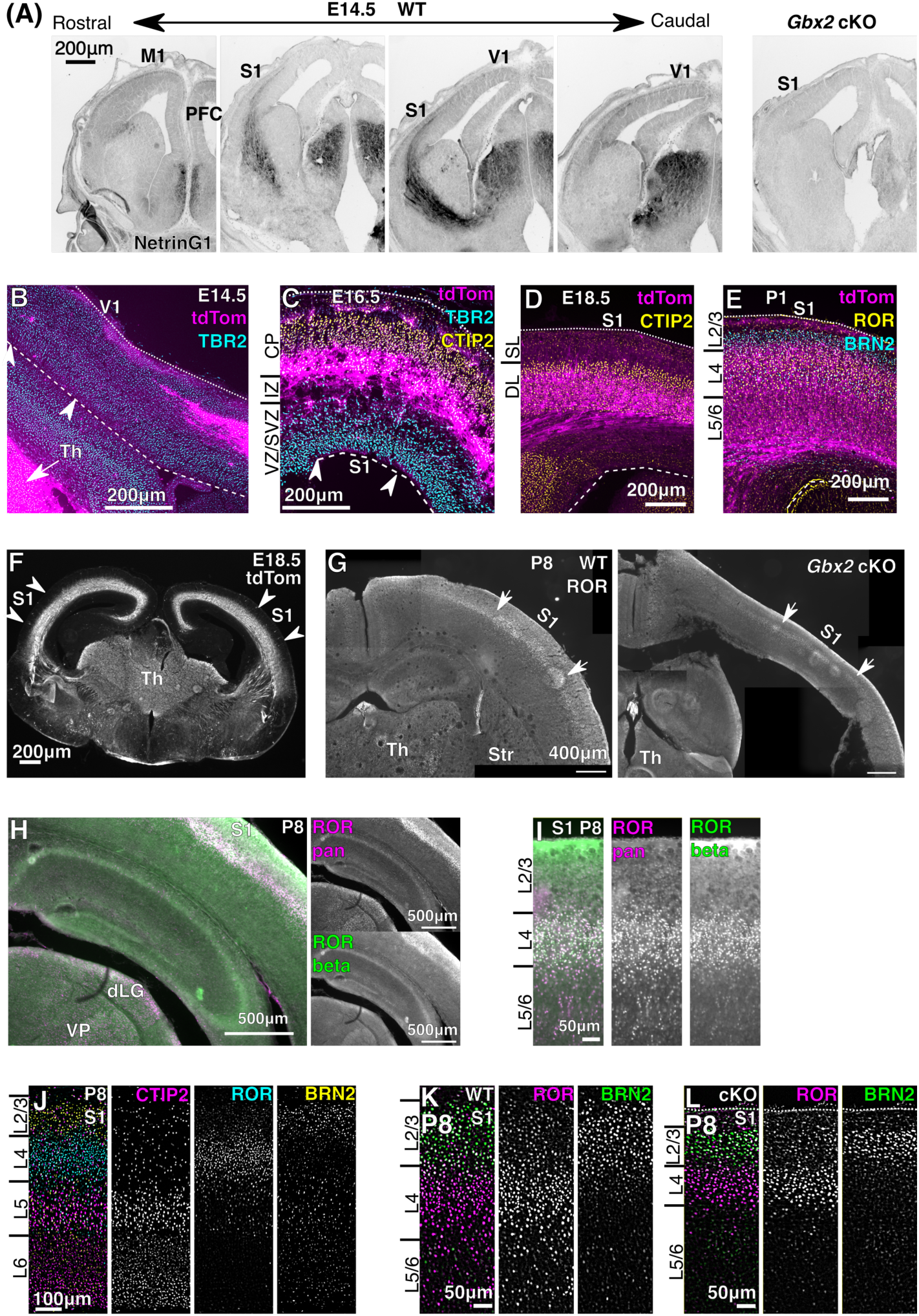
Thalamocortical axons reach the neocortex during neurogenesis, which is prevented in conditional *Gbx2* mutant mice. **(A)** By E14.5, TCAs (NetrinG1; inverted) reach primary somatosensory (S1) and minimally to primary visual (V1) but not to primary motor (M1) or prefrontal cortex (PFC). *Gbx2* cKO coronal section shown to compare to middle S1 section and shows no axons present. **(B-E)** Fluorescent reporter tdTomato driven by *Olig3^Cre^* shows extent of TCAs in embryonic and neonatal stages. At E14.5, TCAs arrive near the lateral part of prospective V1 cortex (B). At E16.5, TCAs reside superficial to intermediate progenitors (TBR2) and below the layer of early-born postmitotic neurons (CTIP2) (C). At E18.5, TCAs have grown up in to the CTIP2+ deep layer neurons, with a few processes beginning to reach above deep layers (D). By P1, TCAs robustly innervate layer 4 (ROR+) but not layer 2/3 (BRN2+)(E). **(F)** At E18.5, TCAs from the thalamus (Th) innervate primary sensory areas (e.g. S1) but also other areas of the embryonic neocortex. **(G)** Layer 4 (ROR^+^) is thinner and less clearly bordered laterally in S1 of *Gbx2* cKO mice. (**H**) Comparison between mouse anti-pan-ROR antibody used in this study (“ROR”) and rabbit anti-RORβ antibody (a gift from Dr. Henk Stunnenberg; no longer available) used as a reference. Anti-pan-ROR is in magenta and anti-RORβ in cyan. The signals for these antibodies match well in the cortex, whereas the thalamus shows more robust labeling with anti-pan-ROR, indicating the dominant presence of RORα in the thalamus (dLG, dorsal lateral geniculate nucleus; VP, ventral posterior nucleus) (Nakagawa and O’Leary, 2003). **(I)** A higher magnification of the S1 cortex comparing the pan-ROR and anti-RORβ antibodies. For analyses, ROR cells below layer 4 were excluded. **(J)** Breakdown of triple immunostaining for CTIP2, ROR and BRN2 at P8. (**K,L**) Comparison of double-immunostaining for ROR and BRN2 at P8 between WT (K) and *Gbx2* cKO (L) S1 cortex.

**Figure S2.**
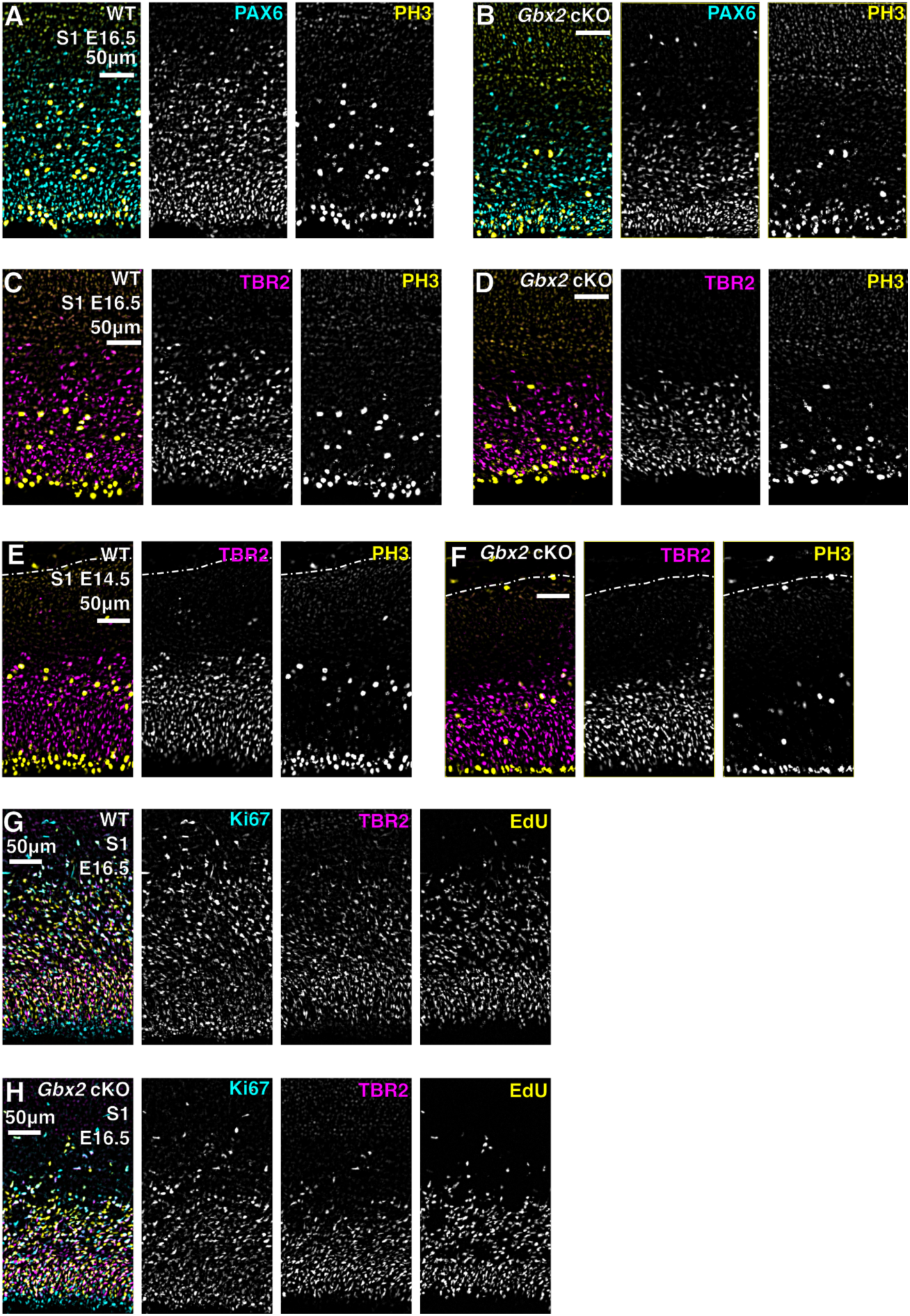
Image breakdowns for progenitors in the embryonic cortex between wildtype and *Gbx2* cKO mice. **(A-B)** Breakdown of immunostaining for PAX6 (radial glia) and PH3 (mitotic cells) at E16.5 for WT (A) and *Gbx2* cKO (B). **(C-D)** Breakdown of TBR2 (intermediate progenitors) and PH3 at E16.5 for WT (C) and *Gbx2* cKO (D). **(E-F)** Breakdown of TBR2 and PH3 at E14.5 for WT (E) and *Gbx2* cKO (F). **(G-H)** Breakdown of triple immunostaining for Ki67 (progenitors), TBR2, and EdU (injected 18-hours prior) at E16.5 for WT (G) and *Gbx2* cKO (H).

**Figure S3.**
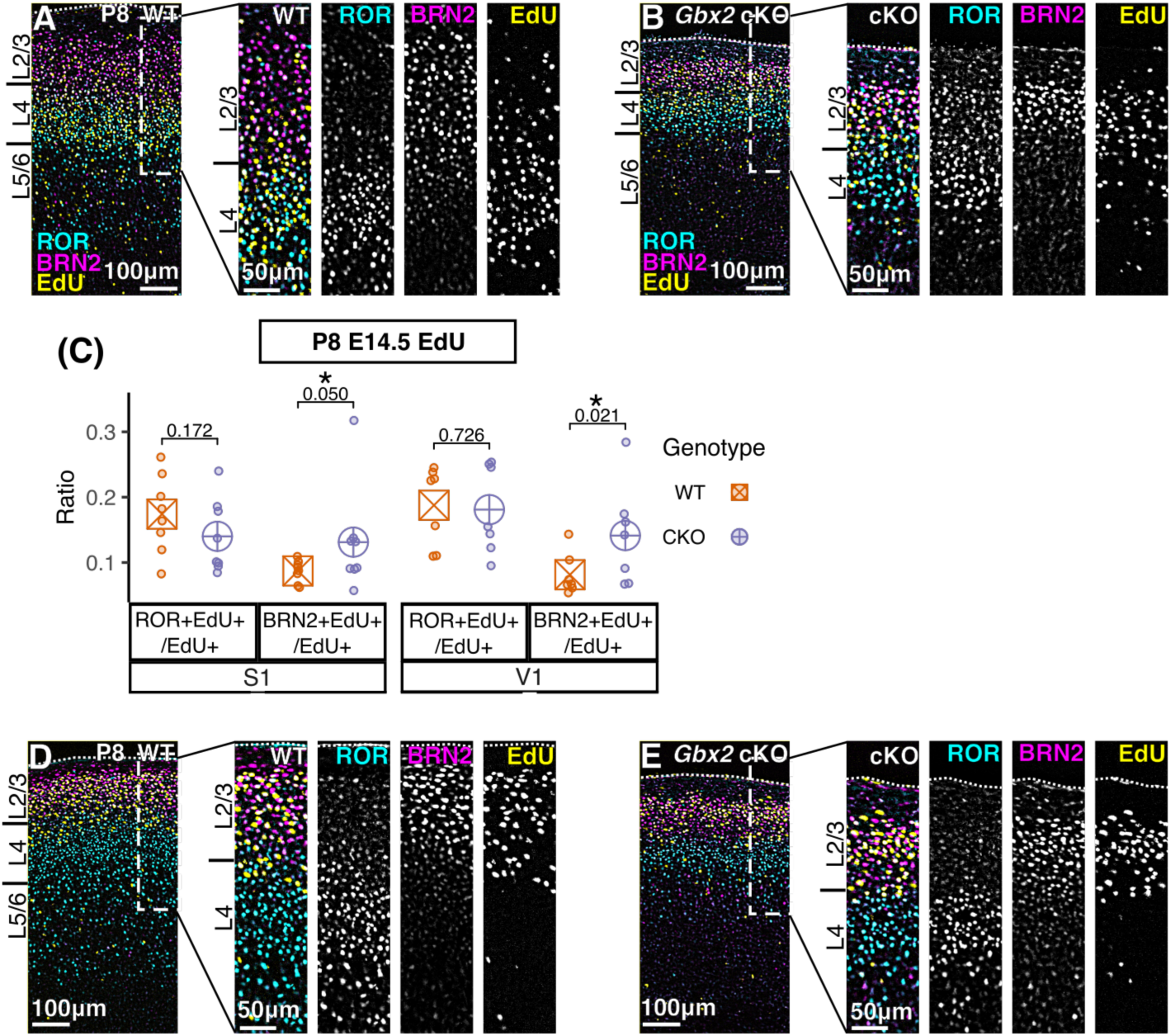
Thalamocortical axons increase layer 4 neurons at the expense of layer 2/3 neurons. **(A-B)** Breakdown of triple immunostaining for ROR, BRN2, and EdU (injected at E14.5) at P8 for WT (A) and *Gbx2* cKO (B). **(C)** At P8, *Gbx2* cKO mice have fewer E14.5-labeled EdU+ cells that acquire layer 4 gene expression (ROR+EdU+/EdU+) in S1 and more cells that acquire layer 2/3 gene expression (BRN2+EdU+/EdU+) in both S1 and V1. **(D-E)** Breakdown of triple immunostaining for ROR, BRN2, and EdU (injected at E16.5) at P8 for WT (D) and *Gbx2* cKO (E). Paired t-test (C) *p*-values shown between WT and *Gbx2* cKO. Circles represent individual brains; large cross-shape represents mean.

**Figure S4.**
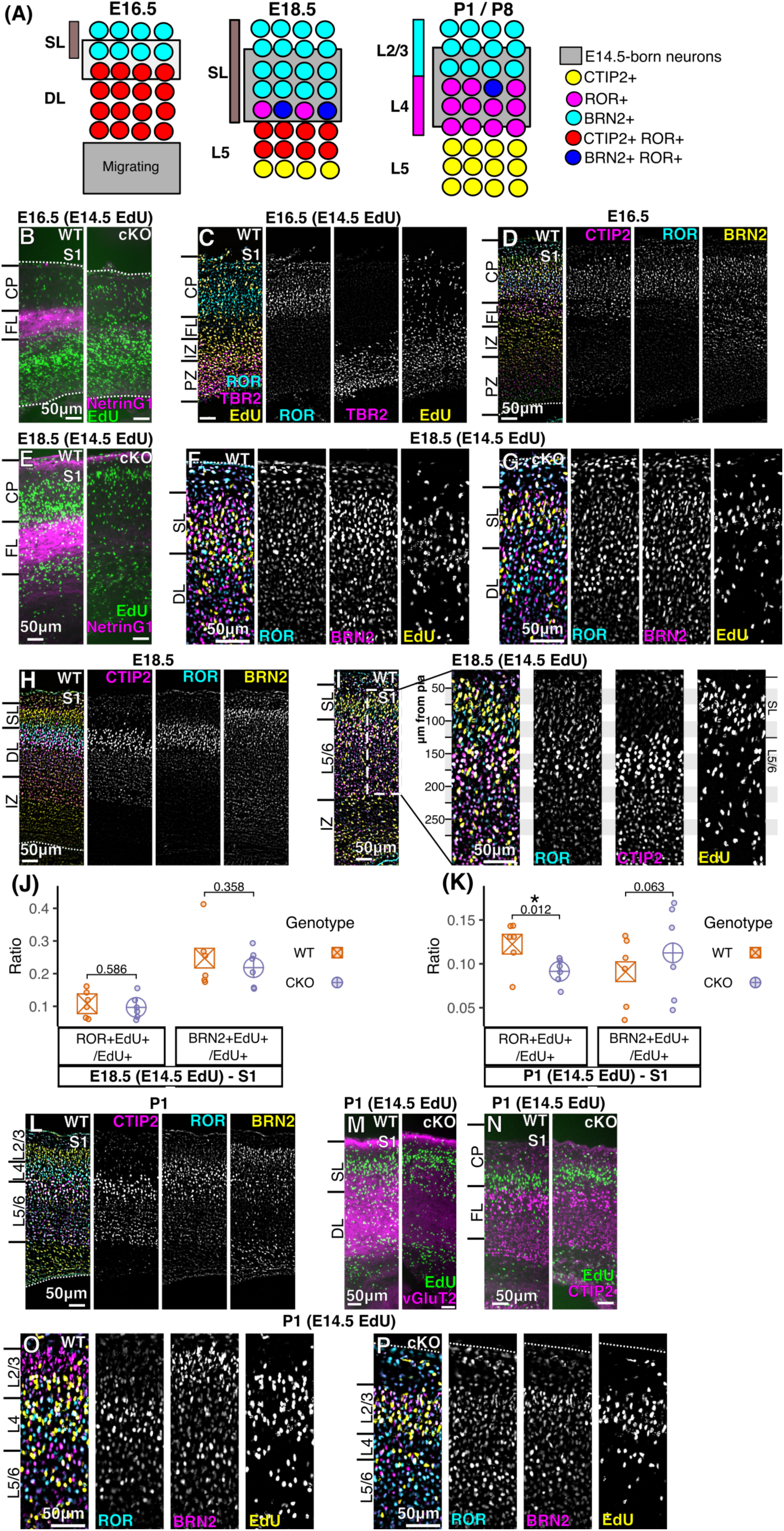
Thalamocortical axons regulate the acquisition of layer 4 gene expression in perinatal sensory cortex. **(A)** Schematic to explain the acquisition of cell fate (L4, ROR+; L2/3, BRN2+) for early born superficial layer (SL) neurons (E14.5-labeled EdU+) across embryonic and neonatal stages in sensory neocortex. At E16.5, EdU+ cells are still migrating through deep layers (DL) with some arriving in the SLs of cortex. At E18.5, BRN2+ cells begin to express ROR and layer 5 (CTIP2) begin to lose expression of ROR. At P1, layer 4 ROR+ cells have acquired differentiated gene expression from both layer 5 (CTIP2) and layers 2/3 (BRN2), which remains consistent through P8. **(B)** Comparison of radial positions of E14.5-born (EdU+) cells relative to TCAs (NetrinG1) at E16.5 between WT (Fig.4B) and *Gbx2* cKO S1 cortex. **(C)** Comparison of ROR+ (neurons in cortical plate), TBR2+ (intermediate progenitors) and E14.5 EdU-labeled cells at E16.5. **(D)** Breakdown of CTIP2, ROR, and BRN2 from Fig.4C in E16.5 S1 WT cortex. **(E)** Comparison of WT (Fig.4B) and *Gbx2* cKO position of E14.5-born (EdU+) cells relative to TCAs (NetrinG1) at E18.5. **(F-G)** Breakdown of ROR, BRN2, and EdU at E18.5 from Fig.4D in S1 of WT (F) and *Gbx2* cKO (G). **(H)** Breakdown of CTIP2, ROR, and BRN2 from Fig.4C in E18.5 S1 WT cortex. **(I)** Comparison of ROR, CTIP2 (layer 5), and E14.5-born (EdU+) cells in WT S1 cortex at E18.5. CTIP2+ cells do not contain EdU at E18.5. **(J)** A higher ratio of E14.5-born (EdU+) cells are BRN2+ compared with ROR+ at E18.5 in both WT and cKO S1 cortex. **(K)** By P1, the ratio of E14.5-born cells is increased for ROR+ cells and decreased for BRN2+ cells when compared with E18.5 in WT mice; in *Gbx2* cKO mice, the ratio of ROR+EdU+ over EdU+ cells is reduced. **(L)** Breakdown of CTIP2, ROR, and BRN2 from Fig.4C in P1 S1 WT cortex. **(M)** Comparison of WT (Fig.4B) and *Gbx2* cKO positions of E14.5-born (EdU) cells relative to TCAs (vGluT2+) at P1. **(N)** E14.5-born (EdU+) cells reside above CTIP2 cells (layer 5) at P1. **(O-P)** Breakdown of ROR, BRN2, and EdU at P1 from Fig.4E in S1 of WT (O) and *Gbx2* cKO (P). Paired t-test (J,K) *p*-values shown between WT and *Gbx2* cKO. Circles represent individual brains; large cross-shape represents mean.

**Figure S5.**
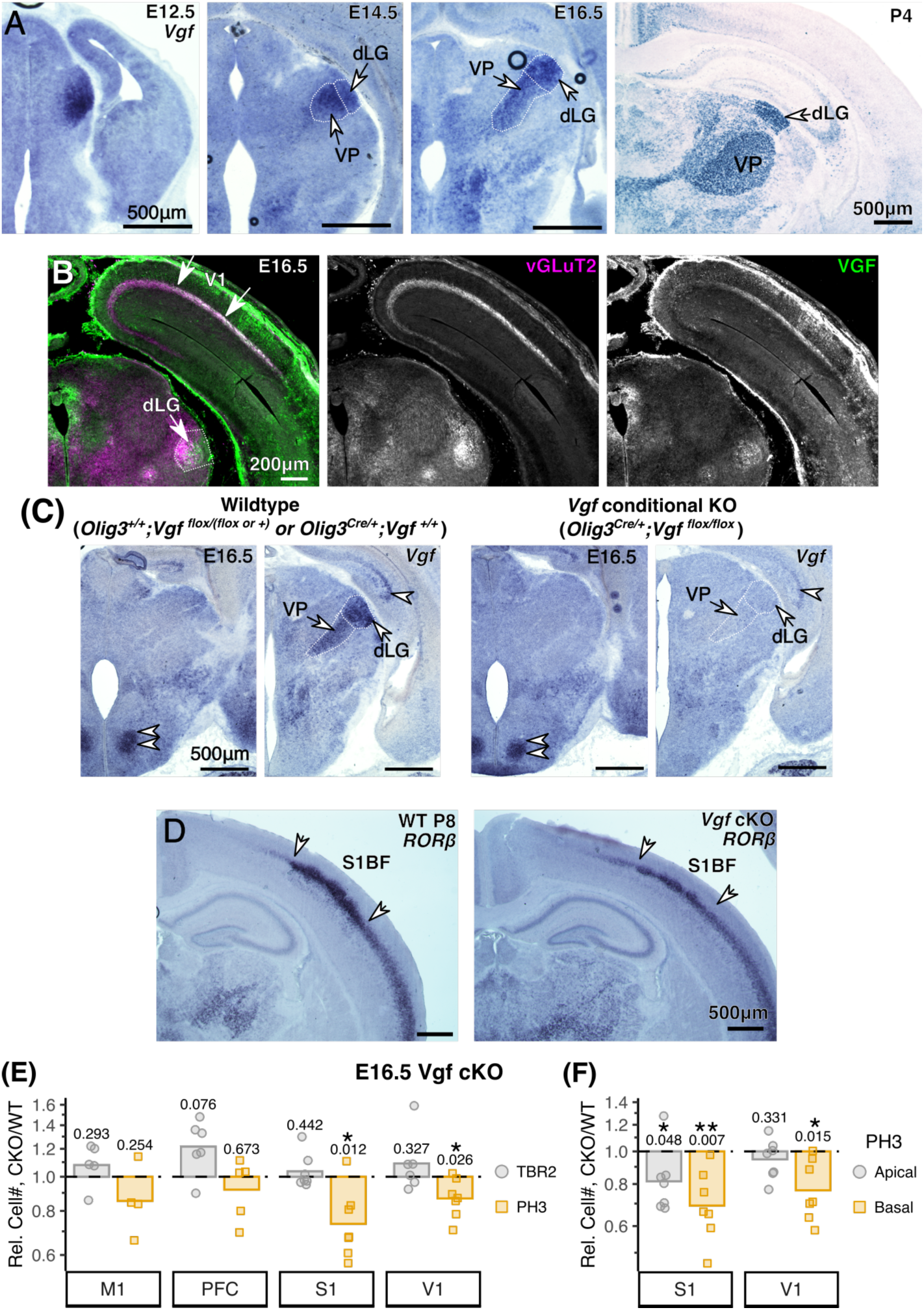
*Vgf* is expressed in principal sensory nuclei of embryonic thalamus and regulates progenitors in the neocortex. **(A)** *In situ* hybridization for the *Vgf* mRNA. *Vgf* mRNA is detected on coronal sections of both embryonic and early postnatal mice. The principal sensory thalamic nuclei, dorsal lateral geniculate (dLG), ventral posterior (VP), and ventral medial geniculate (not shown), all express *Vgf*. The neocortex does not express *Vgf* except in the medial cortex at P4. **(B)** VGF is immunolabeled in the terminals of thalamocortical axons (vGLuT2+) and are also shown here in the dLG in the thalamus. **(C)** Validation of thalamus-specific deletion of *Vgf* mRNA in *Olig3^Cre/+^*; *Vgf^flox/flox^* (*Vgf* cKO) mice. Thalamic expression of *Vgf* (arrows) is specifically lost in the thalamus of the cKO mice, whereas *Vgf* mRNA in the hypothalamus (double arrowhead) and hippocampus (arrowhead) is preserved. **(D)** *Vgf* cKO show a thinning of layer 4 (*RORβ* mRNA) and lack of boundaries between S1 barrel field (S1BF) and adjacent areas (arrows). **(E)** At E16.5, the number of intermediate progenitors (TBR2) is not different between WT and *Vgf* cKO mice in sensory areas S1 and V1. The overall number of mitotic cells (PH3) cells is decreased in both S1 and V1 of *Vgf* cKO mice, but is unchanged in M1 and prefrontal cortex (PFC). **(F)** At E16.5, the number of PH3+ cells in the basal position is decreased in both S1 and V1, whereas PH3+ cells in the apical position are decreased in S1. Matched ratio t-test (E,F) *p*-values shown between WT and *Vgf* cKO. For (E,F), data shown as relative difference of *Vgf* cKO to WT littermates (dashed line); points represent individual pairs, bar represents pooled mean.

